# A hierarchical strategy to decipher protein dynamics *in vivo* with chemical cross-linking mass spectrometry

**DOI:** 10.1101/2023.03.21.533582

**Authors:** Beirong Zhang, Zhou Gong, Lili Zhao, Yuxin An, Hang Gao, Jing Chen, Zhen Liang, Maili Liu, Yukui Zhang, Qun Zhao, Lihua Zhang

## Abstract

Protein dynamics are essential for their various functions. Meanwhile, the intracellular environment would affect protein structural dynamics, especially for the intrinsically disordered proteins (IDPs). Chemical cross-linking mass spectrometry (CXMS) can unbiasedly capture the protein conformation information in cells and can also represent the protein dynamics. Here, we proposed a hierarchy deciphering strategy for protein dynamics *in vivo*. With the prior structure from AlphaFold2, the steady local conformation can be extensively evaluated. On this basis, the full-length structure of multi-domain proteins with various dynamic features can be characterized using CXMS. Furthermore, the complementary strategy with unbiased sampling and distance-constrained sampling enables an objective description of the intrinsic motion of the IDPs. Therefore, the hierarchy strategy we presented herein could help us better understand the molecular mechanisms of protein functions in cells.

## Introduction

Proteins are the main functional macromolecules in biological life processes. Their structure and dynamics are responsible for interaction assembly, driving the function regulation ^[1]^. Until now, a series of experimental techniques have made substantial achievements in protein structural analysis. For example, x-ray diffraction technology has outlined 87.1% of the protein structure in the Protein Data Bank (PDB) using recombinant protein samples expressed *in vitro* ^[2]^. NMR and cryo-electron microscopy have also extensively studied protein structures ^[3]^. However, more than half of the 20,375 gene-coding homo sapiens proteins deposited in the UniProt database (www.uniprot.org) have not been experimentally identified.

The AI system-AlphaFold was developed in recent years to predict the protein tertiary structure from amino acid (AA) sequences ^[4]^. Especially the latest upgraded version, AlphaFold2, significantly improves the prediction accuracy ^[5]^, ushering in a new era in digital biology. The AlphaFold2 has provided over 200 million full-length protein structures with relatively high accuracy, especially for the stable well-folded regions. However, the AlphaFold2 predicted structure is derived from the training dataset depending on the existing structure (or structure fragment) in the PDB database. Limited by the optimization algorithm, the predicted structures by AlphaFold2 were typically single, static conformations lacking dynamic features. Meanwhile, the accuracy for AlphaFold2 is poor for the flexible regions that have not been experimentally resolved (e.g., the missing regions in crystal structures).

The protein dynamics are the essence of their various functions. The protein dynamics were commonly provided by flexible regions, including the intra and inter-loop regions within/between the diverse structural domains, which is essential in folding and conformational transitions. The dynamics of flexible regions dramatically increase the resolving difficulty of the full-length protein structures, resulting in preferences for the relatively rigid parts by current structural biology methods. While the dynamic changes in the flexible regions closely affect the overall conformation, which is essential for function exploration ^[6]^. The intrinsically disordered proteins (IDPs) are typical examples of protein structure dynamics. The IDP molecule lacks a stable tertiary structure, making it difficult to resolve its structure. This also leads to the inability of AlphaFold2 to predict the structure of IDPs accurately. On the other hand, IDPs play various vital functions in cells. For example, many IDPs have been reported to be relevant to the dynamic assembly process of liquid-liquid phase separation and disease states ^[7]^. In addition, the protein dynamics will be affected by the surrounding environment. The complex environment in the cell will cause proteins to exhibit different structural and dynamic properties. As a result, studying protein dynamics, especially the dynamic features of protein *in vivo*, is of great significance for us to deeply understand the functional states of proteins.

Chemical cross-linking coupled with mass spectrometry (CXMS) has been quickly gaining momentum as a powerful tool for the study and elucidation of protein conformation ^[8]^. This technology uses a cross-linker containing two reactive groups separated by the arm-specific length to bridge two residues in proximity. The snapshot of fixed structures is then resolved by cross-link identification based on proteomic analysis. In addition, *in vivo* CXMS profiling allows protein conformations to unravel in their native cellular environments by providing the intra- and inter-protein cross-linked site distance information ^[9]^ (Figure 1A). Interactively, the identified cross-links can be used not only for validating AlphaFold2 predicted structures in the experiment, but also for fine-tuning to light up protein structural dynamics of functional states in living cells based on prior structural information by structural modeling strategy.

**Figure 1.**
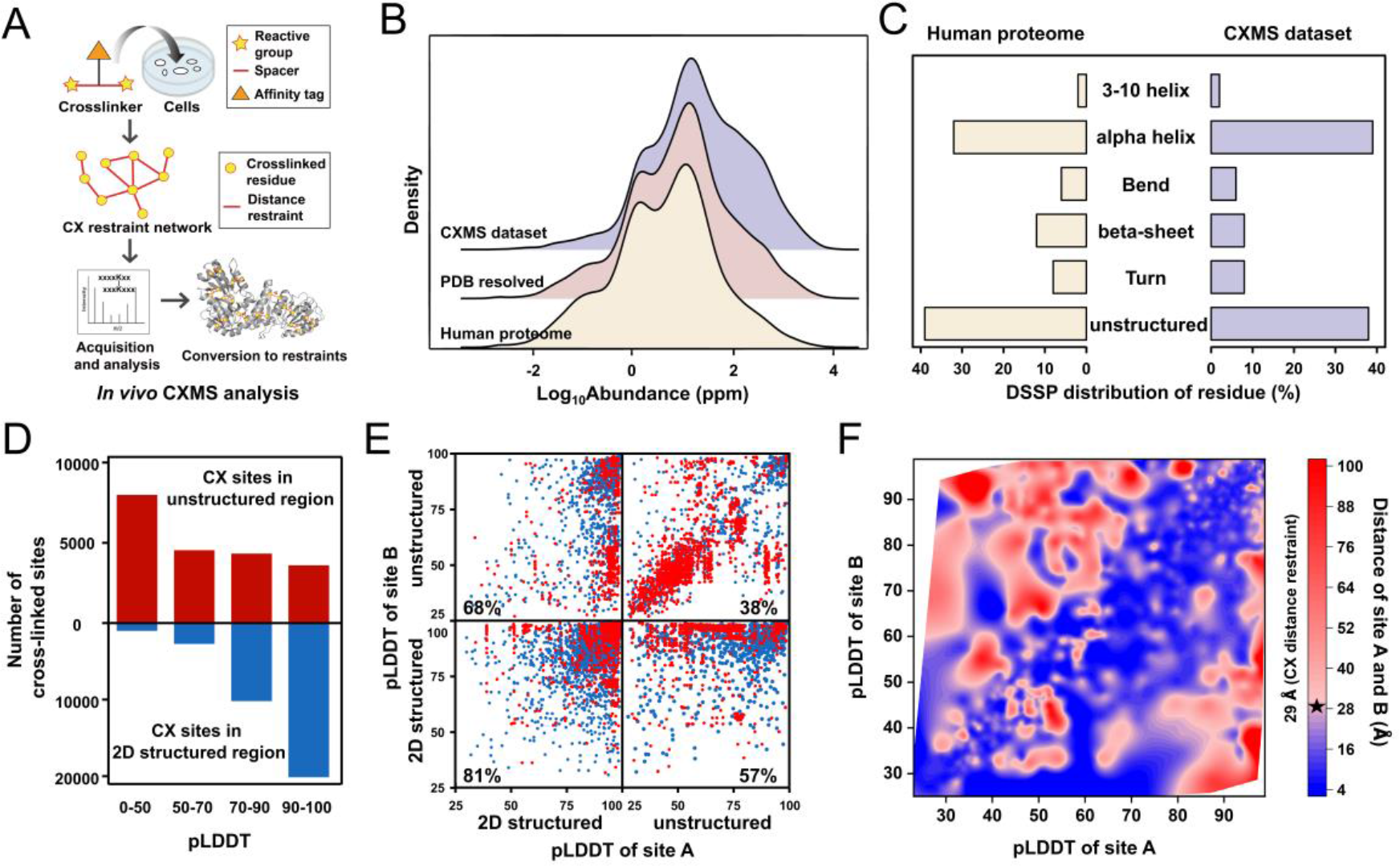
Consistency of cross-linked data for structure deciphering. A) Schematic diagram of CXMS for structural analysis. B) Comparison on abundance distribution of CXMS dataset with human proteome and PDB resolved dataset. C) Comparison on DSSP (dictionary of protein secondary structure) distribution of cross-linked sites in CXMS dataset with residues of human proteome. D) pLDDT score distribution of cross-linked sites, respectively in 2D structured and unstructured regions. E) Structural compatibility of cross-links with AlphaFold2 in 2D structured and unstructured regions. The space number of AA residues between the cross-linked sites were more than 7. Red dot indicated the cross-links incompatible with the AlphaFold2 structure, as the distance calculated between the cross-linked residues exceeding the maximum length of the cross-linker, while blue dot shown compatible. Number in the corner of each category indicated the percentage of compatible cross-links. F) Relevance of structural compatibility of cross-linking with pLDDT score for IDPs. The maximum Cα–Cα distance restraint of cross-linked site pairs was indicated.

In this work, to decipher the protein dynamics in the cell, we have developed a hierarchical strategy to decipher protein dynamics *in vivo* with CXMS. With the prior structure from AlphaFold2, the local domain structure can be evaluated and identified. After that, the dynamic conformation of multi-domain proteins with flexible linkers can be characterized with distance restraints from CXMS. Moreover, the ensemble structure of IDPs was uncovered with restraints molecular dynamics (MD) simulations and unbiased MD simulations, respectively. These two complementary approaches can comprehensively reflect the structural features of IDP in cells. Altogether, these results demonstrated the potential of *in vivo* CXMS to further explore the function of intracellular protein structure and dynamics ^[10]^.

## Results and discussion

### Cross-linking unbiasedly captured diverse protein structures and dynamics in cells

The ability of CXMS to capture reachable sites in different structural regions of proteins directly determines its suitability for structural analysis. In our CXMS dataset previously identified, 26,261 intra-protein cross-links corresponding to 4,914 proteins of human proteome were achieved by *in vivo* CXMS ^[11]^, with respect to 2,690 PDB resolved and 2,224 unresolved structures (Table S1). These proteins are unbiasedly involved in the whole abundance range (Figure 1B) and distributed in all the cellular compartments with diverse molecular functions and sequence lengths (Figure S1A, 1B). Especially the proteins not deposited in PDB were also universally included in CXMS dataset. Therefore, a valuable potential is existed for cross-linking in structure deciphering for various proteins in the cells.

In addition, the full-length predicted protein structures from the AlphaFold2 database were used to investigate the structural compatibility of CXMS dataset. Because of the local flexibility of protein residues, Cα-Cα distances of lysine site pairs within 29 Å were used as the criteria to evaluate the compatibility ^[12]^ (Figure S2). Since the distance between two adjacent residues on the peptide chain was about 4 Å, the two AAs within the spacer seven can always be cross-linked regardless of the protein structure. The distribution of cross-linking sites on the protein indicated that these adjacent AAs were indeed more likely to be cross-linked. In contrast, about 42% of cross-linked AAs have a sequence separation of more than 7 residues. Among them, 25 % of cross-linked residues were separated by more than 20 AAs. These long-range cross-links could contribute important information in describing the protein conformation and dynamics (Figure S1C).

On the other hand, the cross-linked sites were contained in various structural regions (Figure 1C). Importantly, 68% of sites were contained in 2D structured elements, while the others were in unstructured regions. Low predicted local-distance difference test (pLDDT) scores were assigned for the sites included in unstructured regions (Figure 1D). Meanwhile, by matching with AlphaFold2 structure, the two cross-linked sites located both in the unstructured regions had the lowest structural compatibility of 38%. In contrast, those cross-linked sites both in 2D structured were with good structural satisfaction of 81% consistency (Figure 1E). By mapping the cross-linking sites onto a well-folded structure, the structural compatibility could achieve nearly 100% (Figure 2A). As the length of unstructured flexible regions increased, the protein structure became more dynamic and less compatible with cross-linking data. For 60S ribosomal protein L23a, up to 7 cross-links were mismatched. These cross-links located in unstructured flexible beta-like import receptor binding (BIB) domain (residues 32-74). The flexible N-terminal domain of L23a is responsible for the recruitment of nucleic acids for the ribosome binding, thus the mismatched distances might indicate the various functional states or the assembly process of ribosome captured by *in vivo* CXMS (Figure 2B).

**Figure 2.**
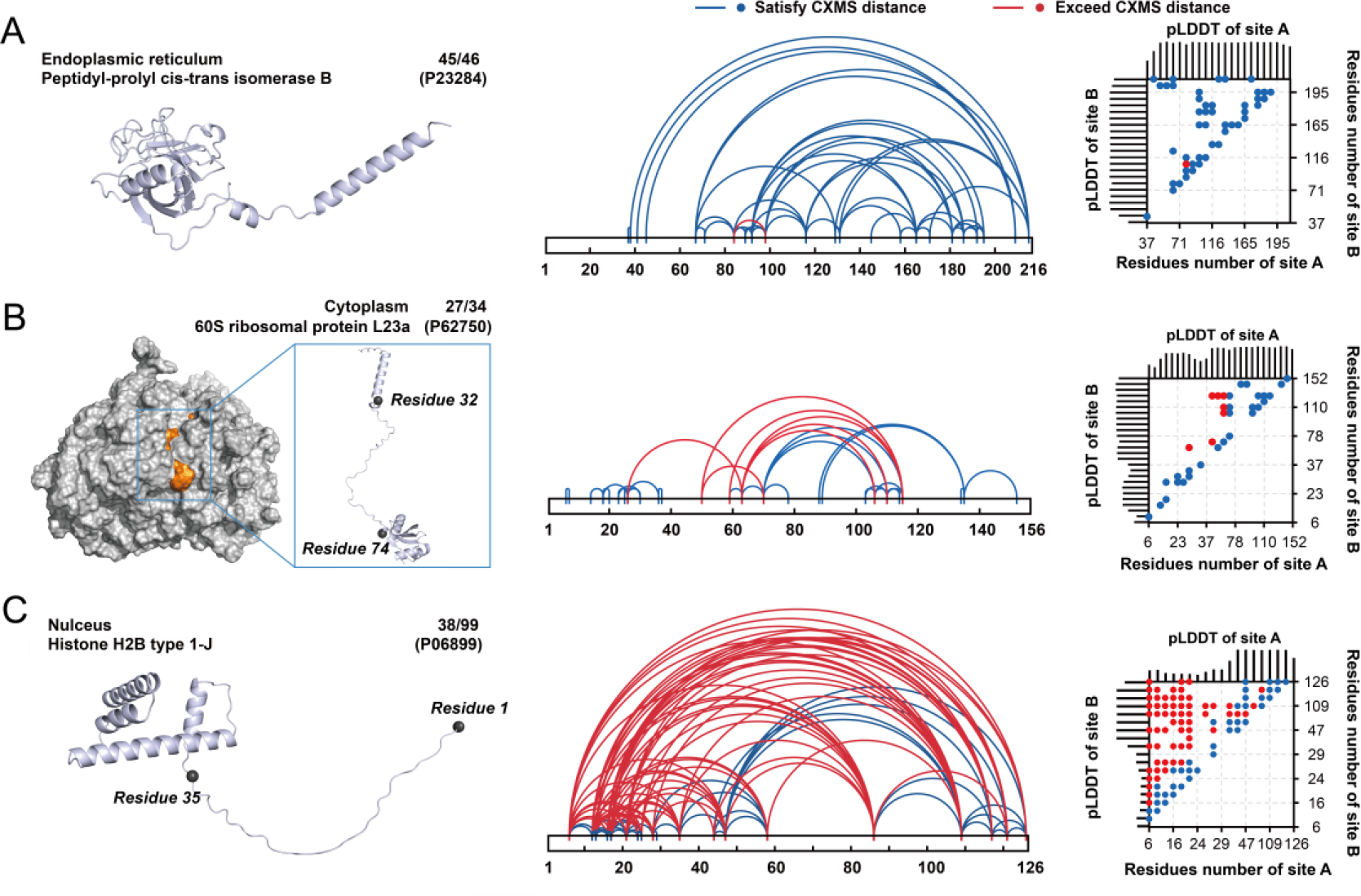
The dynamic conformation characterization for proteins with increasingly unstructured regions by CXMS. The dynamic conformation characterization for peptidyl-prolyl cis-trans isomerase B (PPIB) (A), 60S ribosomal protein L23a (B), and histone H2B (C). In each of these diagrams, left: the AlphaFold2 structure. The number of cross-links within the distance and the total number of cross-linked pairs were indicated and separated by “/”. Middle: all identified cross-links were mapped onto the 2D structure. For the 2D maps, the distance of corresponding residues that satisfy or exceed cross-linking information were colored in blue and red, respectively. Right: correlation between the pLDDT score of cross-linked sites and the structural compatibility of cross-links.

Furthermore, we moved on to inspect the IDPs, which is regarded highly dynamic with lots of unstructured regions. The results indicated the distance between the corresponding sites of AlphaFold2 predicted IDP structure often far exceeded the maximum arm length of the cross-linking agent (Figure 1F). Taking histone H2B as example, except for the adjacent AAs, the distance of most of the sites were mismatch with the cross-linking data (Figure 2C). Meanwhile, due to the highly dynamics of the IDP structure, the prediction accuracy of many sites in the AlphaFold2 predicted structure were low. In addition, these flexible, unstructured regions (especially for IDPs) were more susceptible to the cellular environment. The *in vivo* CXMS can unbiasedly obtain distance information between AAs of different protein types and regions. Therefore, it is critical and reasonable to characterize the different conformational states of these flexible proteins in the cell using *in vivo* CXMS.

### Hierarchical strategy for deciphering protein dynamics with CXMS

Characterizing protein structure and dynamics based on CXMS is essentially to refine the protein structure based on distance restraints. The NMR typically acquires thousands of distance restraints for the structure calculation. In contrast, CXMS can usually only obtain tens or even less cross-linked sites information. As a result, it is particularly important to effectively use some prior structural information for the CXMS calculation. Here, we developed the hierarchical strategy to study the protein dynamics with the prior structure and distance restraints from CXMS. We designed and applied the strategy to characterize the two representative types of proteins, multi-domain proteins and IDPs.

We first analyze the dynamics of multi-domain proteins. This group of proteins are ubiquitous in cells, and usually connected by flexible loop regions. The conformational changes of these flexible loops determine the dynamics of proteins, but make it difficult to resolve the full-length structure, thus only a certain stable conformational state can be resolved. Meanwhile, this manifests an incompatible between these resolved structures and the cross-linking information, i.e., the distance between the corresponding sites was too long relative to the maximum arm length of the cross-linker. This so-called “over-length” cross-linking information can provide the dynamic features of multi-domain proteins (Figure 3A). Protein domains often have independent and stable tertiary structures that can independently or cooperatively perform biological functions ^[13]^. These domain conformations with stable tertiary structure are usually easy to resolve or can be accurately predicted by AlphaFold2. In addition, the intra-domain CXMS information can also be used to verify the domain structures. Therefore, we can perform rigid-body refinement with the prior domain structures and inter-domain CXMS information. In the rigid body refinement process, the domain was considered as a whole, focusing on the spatial position relationship between the domains, which was restrained by inter-domain distance from CXMS. When all the cross-linking restraints between domains were satisfied, we could obtain the ensemble conformation of multi-domain proteins.

**Figure 3.**
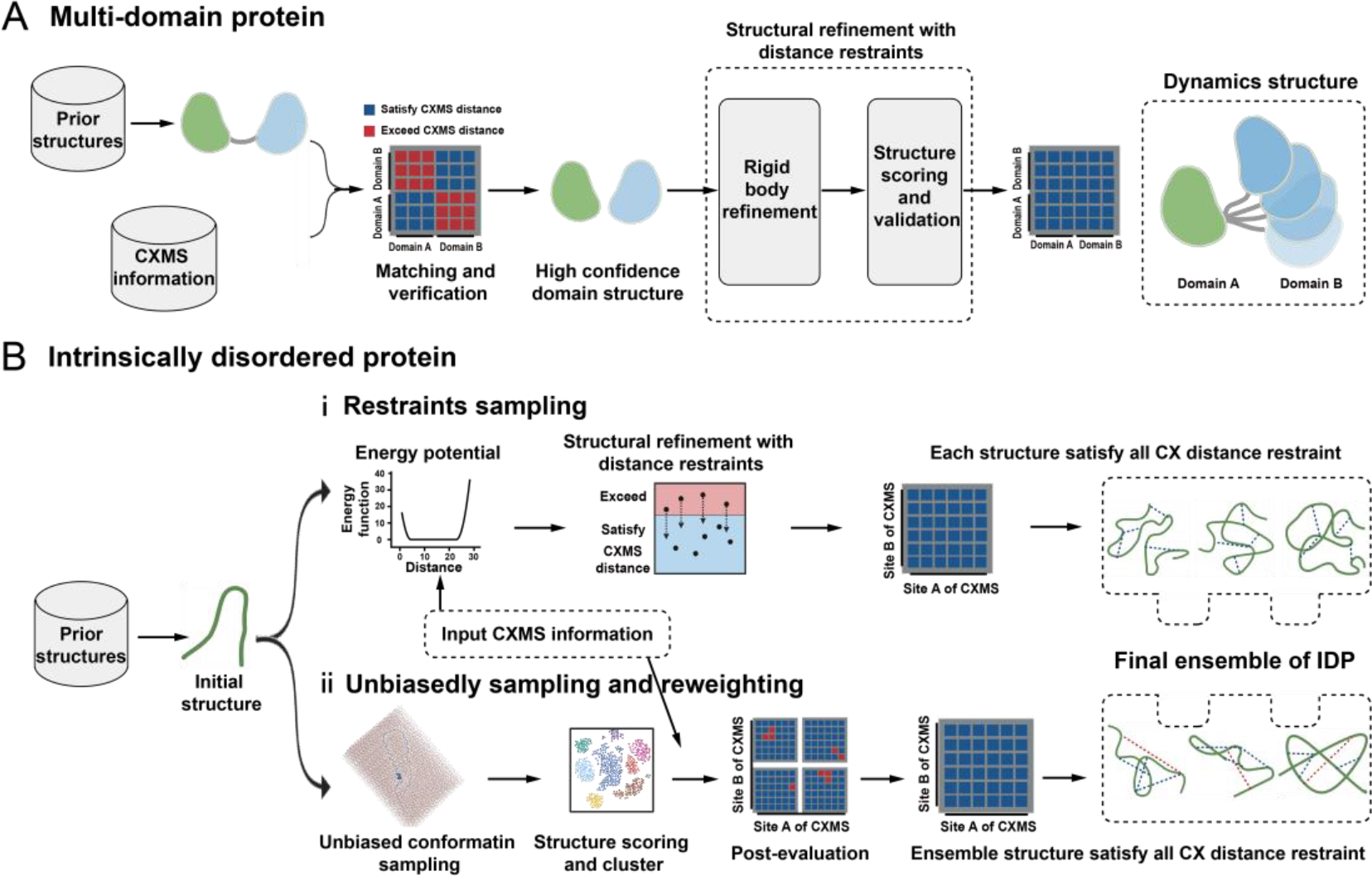
Algorithm workflow for hierarchical deciphering protein dynamics with CXMS. A) The workflow of deciphering ensemble conformation of multi-domain proteins. B) The strategies for deciphering ensemble conformation of IDP. (ⅰ): the structure calculation with distance restraints from CXMS. (ⅱ): unbiasedly sampling combining with post-evaluation by CXMS. Arrows represented the information flow among the different element in the strategy. The grid indicated the matching of cross-linking information.

On the other hand, compared with multi-domain proteins, there is little reliable prior structural information for IDP molecules. The highly dynamic nature of the IDP molecule gave us more freedom in choosing the initial structure for the structure calculation. The initial structure of IDP can be obtained from the PDB database, AlphaFold2 prediction, or other algorithms like Flexible-meccano ^[14]^. There were two different strategies to obtain the ensemble structure of IDPs combining with CXMS (Figure 3B). At first, the structural refinement with distance restraints from CXMS can be performed with the same process as the multi-domain proteins. Importantly, the main difference was that there was no rigid body part for the IDP molecule. The entire protein was freely sampled during the structure sampling process. The final calculated structures satisfied all the distance restraints would be selected for the further analysis (Figure 3B-ⅰ). The advantage of this algorithm was the high sampling efficiency.

However, since the lack of stable tertiary structure of IDP, too many distance restraints in the IDP sampling process might bring artificial errors for the final calculated ensemble conformations. Therefore, the ensemble structure obtained by this restraint approach required further validation.

As an alternative, the unbiasedly sampling combining with post-evaluation can objectively obtain the intrinsic motion state of IDP. The distance information from CXMS can further reweight the sampling structure. Totally, the IDP was unbiasedly sampled with all atom MD simulations to produce the large conformation pool. The conformations satisfying some or all the distance restraints were selected. In addition, the obtained structures were further analyzed using clustering or energy evaluations (Figure 3B- ⅱ). The very important advantage of this strategy was the unbiased sampling enabling the results more objective. But ensuring the adequacy of sampling was needed to make the results reliable. Altogether, the structure calculation with CXMS information should make best use of prior structural information and distance restraints as much as possible, whether as a guidance during the sampling process or as a re-evaluation or reweighting after sampling. The structures obtained by these different approaches can complement each other and confirm each other.

Then, as a proof of concept, we will demonstrate the exemplification of the above hierarchical strategies for representation of inter-domain dynamics, as well as the ensemble conformation of IDPs.

### Inter-domain dynamics captured by *in vivo* CXMS

First, we visualized the inter-domain dynamics of calmodulin (CaM, UniProt ID: P0DP23), which performs regulatory functions through calcium binding. The CaM protein has a dumbbell-like extended conformation without ligand binding. The N-terminal domain (NTD) and C-terminal domain (CTD) of AlphaFold2 predicted structure with a high pLDDT score agrees with the *apo* crystal structure (PDB: 1CLL). The residues between NTD and CTD (residues 77-81) have a low pLDDT score (< 60), leading to different overall structures (Figure 4A and Figure S3A). Therefore, the dumbbell structure can be bent and twisted due to the flexibility of the middle residues. Five *in vivo* cross-links of CaM were identified with two inter-domain cross-links (K14-K95 and K22-K95) (Table S2). The distance between the inter-domain cross-linked residues in the AlphaFold2 predicted structures were 39.0 Å and 47.1 Å, respectively (Figure 4B). These distances far exceeded the maximum arm length of the cross-linker. Thus, a closed state with a shorter distance between these residues could exist in the cell. As such, we performed structure refinement with the distance restraints from CXMS. The NTD and CTD domains were treated as rigid bodies connected by a flexible linker. The alternative closed conformation of CaM was identified from the CXMS restraints. Interestingly, the closed conformation calculated with CXMS was similar to the crystal structure of the ligand-bound *holo* state of CaM (PDB: 2BE6) (Figure 4C). As a result, this close state might come from the specific *holo* state with different ligands bound in the cell. The results also indicated that the closed state could exist without ligand binding, suggesting a conformational selection mechanism in the cell, that was also reported in the *in vitro* studies ^[15]^.

**Figure 4.**
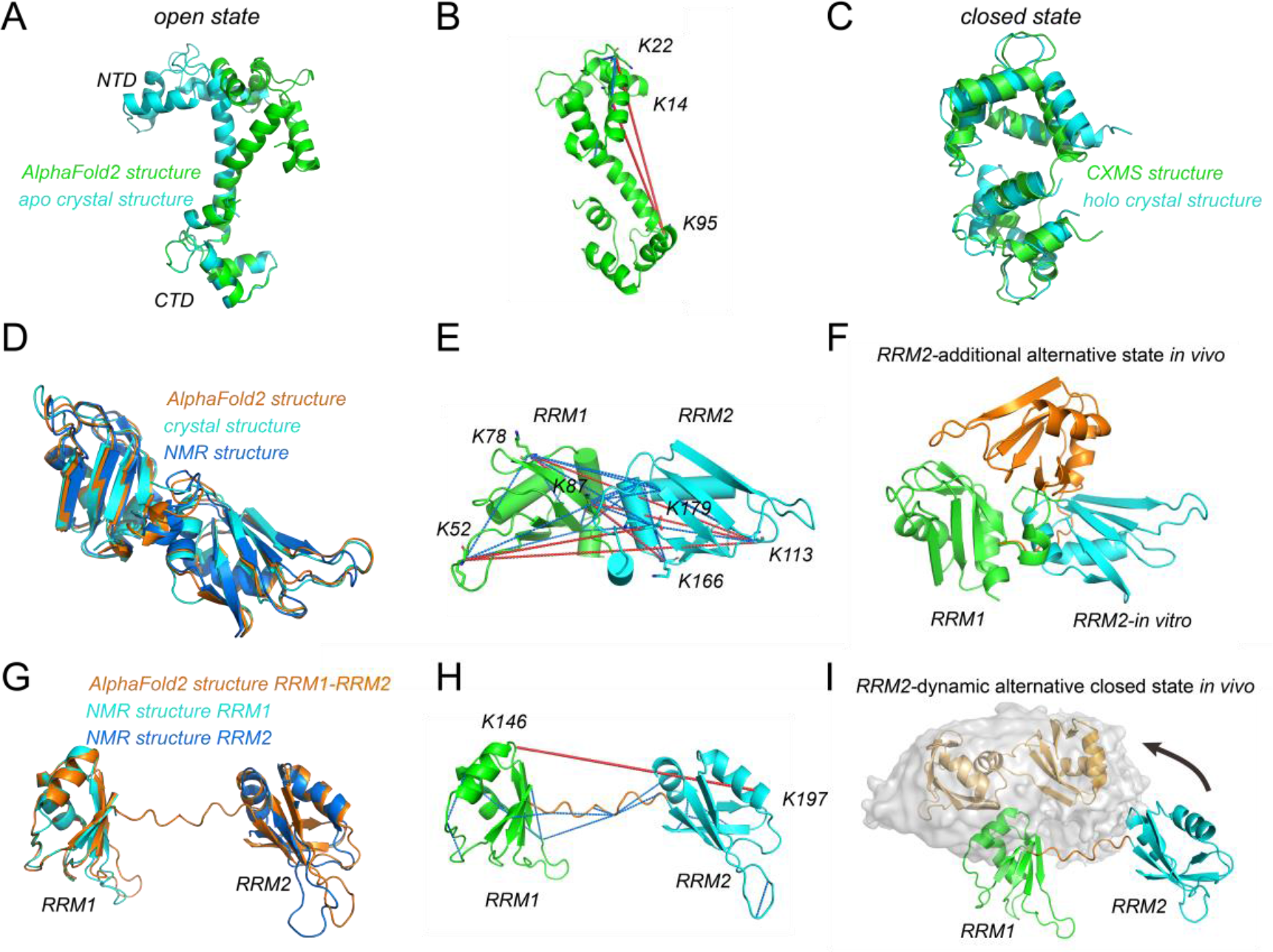
Inter-domain dynamics captured by CXMS restraints. A) Comparison of AlphaFold2 and crystal structure (PDB: 1CLL) of CaM (UniProt ID: P0DP23). Both two structures were in open conformation state. B) Mapping all cross-links onto the AlphaFold2 predicted structure of CaM. C) Structural comparison of calculated structure with CXMS restraints and crystal structure of ligand-bound CaM (PDB: 2BE6). Both two structures were in closed conformation state, and the RMSD was 3.95 Å for all heavy atoms. D) Comparison of AlphaFold2, NMR (PDB: 2LYV), and crystal structure (PDB: 1HA1) of RRM domains of hnRNP A1(UniProt ID: P09651). E) Mapping all cross-links onto the AlphaFold2 predicted structure of hnRNP A1 RRM domains. The RRM1 and RRM2 domains were expressed in different colors. F) Additional alternative structure of RRM2 (orange) of hnRNP A1 with respect to RRM1 in the ensemble conformation refined against CXMS data. For comparison, the AlphaFold2 structure was superimposed on a cartoon. G) Comparison of RRM1-RRM2 structure of hnRNP D0 (UniProt ID: Q14103) predicted by AlphaFold2 and NMR structures for RRM1 (PDB:1HD0) and RRM2 (PDB: 1IQT) respectively. H) Mapping all cross-links onto the AlphaFold2 predicted structure of hnRNP D0 RRM domains. The RRM1, RRM2 domain, and flexible linker were expressed in different colors. I) The surface indicated the distribution of RRM2 of hnRNP D0 with respect to RRM1. The two represented conformation of RRM2 was shown in wheat cartoon. The AlphaFold2 predicted structure was also represented as cartoon. In the B, E and H, cross-links exceeding the maximum distance of cross-linker arm length were represented as red dotted lines, while the others were drawn as blue.

Then, the inter-domain dynamics were also analyzed on the heterogeneous nuclear ribonucleoprotein A1 (hnRNP A1, UniProt ID: P09651), a member of the hnRNP family involved in DNA/RNA binding and RNA splicing. The human hnRNP A1 has 372 AAs, including two RNA recognition motifs (RRMs) at the N-terminus and a low-complexity region (LCD) at the C-terminus (Figure S3B) ^[16]^. The well-folded RRM (RRM1: residues 10-88 and RRM2: residues 106-179) domains have a high pLDDT score (>90 for most residues), while the flexible LCD regions have low prediction confidence. The RRMs structures predicted by AlphaFold2 were consistent with structures resolved by solution NMR or X-ray crystallography ^[17]^. The RMSD between these structures was less than 2 Å for all heavy atoms, indicating that the RRM domains were mainly in a single and stable conformation state *in vitro* (Figure 4D). A total of 54 *in vivo* cross-links were identified for hnRNP A1 with 10 intra-domain cross-links, 11 inter-domain cross-links (Table S3), and 33 cross-links with at least one residue located in the flexible region. Of them, 5 inter-domain cross-links exceeded the maximum arm length of the cross-linker (Figure 4E). These “over-length” cross-linked residues suggested the inter-domain dynamics between two RRMs, resulting in the alternative conformation that differs from the conformation resolved *in vitro*.

Based on the CXMS restraints, the ensemble conformation refinement was performed to characterize the inter-domain dynamics of hnRNP A1 RRMs *in vivo*. The intra-domain cross-links were consistent with the structure in the database, indicating the structure of a single RRM domain should keep stable *in vivo*. As such, we mainly focused on changes in the spatial position between the two RRM domains. The results indicated that an ensemble consisting of two conformers could satisfy all CXMS restraints. One of them was consistent with the experimental resolved structure, while the RRM2 in another alternative conformation moved to the other side of the RRM1 (Figure 4F). As a result, in addition to the structure resolved *in vitro*, the RRM domains would form additional alternative conformational states *in vivo*, which could facilitate RNA recognition and binding.

Furthermore, another heterogeneous nuclear ribonucleoprotein (hnRNP D0, UniProt ID: Q14103) was analyzed with *in vivo* CXMS restraints. The hnRNP D0 also has two RRM domains (RRM1: residues 98-171 and RRM2: residues 183-256) besides two LCD regions at both the N-terminus and C-terminus (Figure S3C). The structures of the two RRM domains have been separately resolved ^[18]^, but the full-length structure containing both RRM domains have not been experimentally determined (Figure 4G). The AlphaFold2 predicted full-length conformation assumes an open conformation, with two RRM domains were separated. In this study, 24 pairs of cross-links were identified for hnRNP D0 *in vivo* (Table S4). The distance between the two cross-linked residues in two RRMS (K146-K197) was 50 Å in the AlphaFold2 structure (Figure 4H). Since the AlphaFold2 predicted structures of these two RRM domains were consistent with the experimentally resolved structures (Figure 4G), we treated the two RRM domains as rigid bodies during the structure refinement. The calculated structures confirmed that the two RRM domains could form closed conformations *in vivo*, that the RRM2 could move around the RRM1 (Figure 4I). These two RRM domains were reported independently bind nucleic acids in the previous studies ^[19]^. Therefore, the closed conformation might enable the two RRM domains to cooperate in nucleic acid binding, resulting in stronger binding capacity or higher selectivity *in vivo*.

Taken together, starting with the prior domain structure from database, we have identified the inter-domain dynamics of three different proteins in cells by structure refinement with the distance restraints from *in vivo* CXMS. Some of these alternative structures have been identified *in vitro* (like closed conformation of CaM), while others have not been reported yet. Furthermore, we also characterized the protein for which the full-length structure was not resolved *in vitro*. Although the AlphaFold2 can predict the full-length structure of these proteins, its overall structure cannot accurately reflect the dynamic properties of the protein due to the low prediction confidence of the loop region. Based on the distance restraints obtained from CXMS, the protein dynamics we have obtained were more accurate and reliable. Besides, these alternative conformations formed due to the influence of the intracellular environment could greatly aid in the discovery of additional protein functional states.

### Deciphering of ensemble conformation of IDPs with *in vivo* CXMS

Due to the extreme flexibility, the IDP structures are always difficult to be resolved. In this study, we refined the ensemble structure of the high mobility group protein HMG-I/Y (UniProt ID: P17096) using *in vivo* CXMS restraints. The HMG-I/Y bind preferentially to the minor groove of A+T-rich regions in double-stranded DNA. Part of this protein (about 20 residues) bound with DNA has been resolved using NMR experiments, but the full-length structure still needs to be determined. The AlphaFold2 predicted structure exhibits a random coil state without any tertiary or secondary structure (Figure 5A). The pLDDT scores for most residues are less than 70, suggesting a low prediction accuracy. A total of 27 *in vivo* cross-linking pairs were identified, and 21 came from the long-range residues (Figure 5B, Table S5). Here, we performed the hierarchical sampling and evaluation strategy to characterize the ensemble structure of HMG-I/Y. First, by performing the structure calculation with distance restraints from CXMS, the ensemble conformation was presented. The conformations that satisfy all distance restraints were mainly distributed in the range of gyration of radius (Rg) 20-30 Å, and end-to-end distance (D_ee_) of 40-80 Å. The top five conformations with lowest energy scores were indicated (Figure S4). Unlike the extended structure predicted by AlphaFold2, these calculated structures exhibited a relatively more compact conformation.

**Figure 5.**
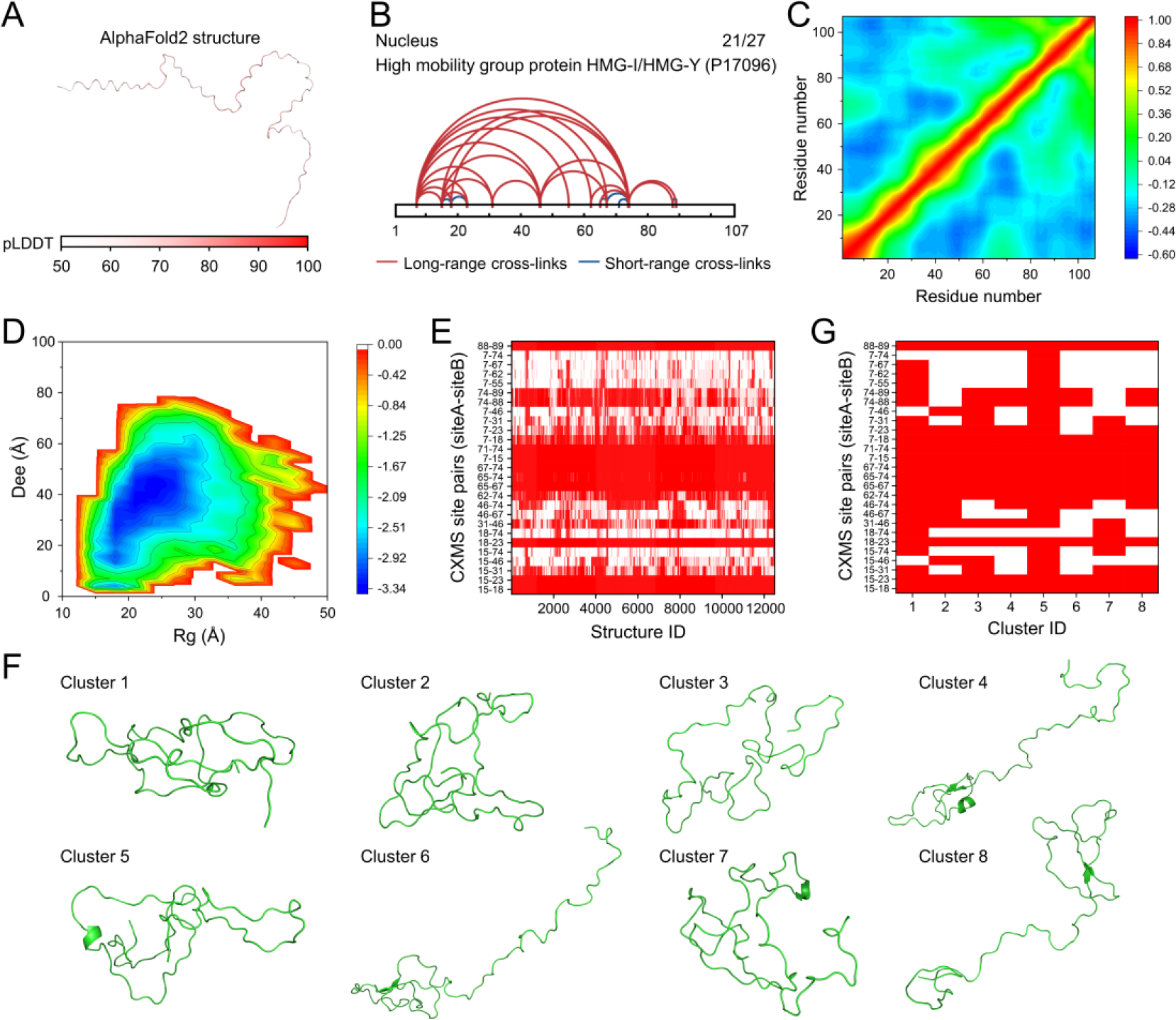
Deciphering of ensemble structure of HMG-I/Y by the strategy of unbiasedly sampling and reweighting. A) The full-length structure of HMG-I/Y predicted by AlphaFold2. The pLDDT scores of residues were indicated by colors from white to red. B) All identified cross-links were mapped onto the 2D structure of HMG-I/Y. The short- (number of residues between cross-linked sites less than 8) and long-range cross-linking information were colored in blue and red, respectively. The number of long-range cross-links and the total number of cross-linked pairs were indicated and separated by “/”. C) The DCCM analysis of HMG-I/Y. The correlation coefficient related with residue related movement were colored from blue to red. The stronger the negative correlation, the small the value of the corresponding position. D) The 2D statistical heat map of Rg and D_ee_ for the MD simulation trajectories of HMG-I/Y. The more the number of structures, the small the value of the corresponding position. E) The matching between each snapshot from MD simulation and corresponding CXMS information. The cross-linking distance satisfying or not was shown in red and white, respectively. F) The representative structure of eight clusters from unbiased all atom MD simulations. G) The matching between eight structure clusters and corresponding CXMS information. The cross-linking distance satisfying or not was shown in red and white, respectively.

In addition, another strategy of the unbiased all atom MD simulations were performed. In order to ensure the sampling sufficiency, the Flexible-meccano algorithm was used to generate the different initial conformations (Rg ranging from 20∼35 Å) for the MD simulations. The dynamical cross-correlation matrix (DCCM) was employed to analyze the intrinsic motion of HMG-I/Y. The results indicated that some residues on the IDP molecule had obvious negative correlation movement, including 15-20 and 40-70, N-terminal and 30-60, 80-100, as well as 30-50 and 60-80 (Figure 5C). This negatively correlated movement suggested a relative motion (like open/close movement) of the whole molecule. Interestingly, these regions with negatively correlated movement were consistent with the cross-linked sites. These results showed that the IDP has an intrinsical compact conformation state that was captured by *in vivo* CXMS. The 2D heat map of Rg and D_ee_ for the MD simulation trajectories showed that the unbiased sampling could obtain a more widely distributed conformations with Rg of 15-35 Å and D_ee_ of 10-60 Å (Figure 5D). We further gained the matching of different cross-linked pairs residues on all the snapshots from MD simulations. It could be found those cross-links between adjacent residues were always satisfied. In contrast, the long-range cross-links could only be satisfied by partial conformations, especially for the cross-links from N-terminal and C-terminal residues (like 7-74, 15-74). Meanwhile, any cross-linking distance could be satisfied on a certain structure. No cross-link pair was always unsatisfactory (Figure 5E). Thus, these results showed the rationality and accuracy of the cross-linking results for characterize the IDP structure in cell. On the other hand, it also indicated that this unbiased sampling could cover the conformation landscape of IDP molecule. In addition, we further clustered the structures from MD simulations. Due to the high dynamic features and high structural heterogeneity of IDP, only 12% of the snapshots were clustered into 8 different classes (Figure 5F). These clustered structures also exhibited a certain compact conformation. Through the analysis of the cross-linking satisfaction of these cluster structures, it was found that these structures could represent the structural properties of the entire simulation trajectories. The ensemble conformation composed of these 8 clusters could satisfy all cross-linking pairs (Figure 5G).

Besides, this analysis algorithm was further applied to other IDP molecules. The non-histone chromosomal protein HMG-17 (UniProt ID: P05204) is a small high mobility group protein lacking a stable tertiary structure. The structure of HMG-17 has not yet been determined by experimental methods. Similarly, the AlphaFold2 predicted structure exhibits a random coil state with no tertiary structures (Figure S5A). We have identified 36 *in vivo* cross-linking pairs, and 17 of them were from the long-range residues (Figure S5B, Table S6). We have performed unbiased sampling strategy to characterize the ensemble structure of HMG-17. The unbiased MD simulation also indicated the negative correlation motions of residues around cross-linked sites (Figure S5C). This indicates that cross-linking objectively captured the dynamic features of this IDP. The conformations of HMG-17 were distributed mainly around Rg of 20-30 Å and D_ee_ of 20-60 Å (Figure S5D). The snapshots from MD simulations could be analyzed into 7 clusters, and most of them exhibits a partially closed conformational state (Figure S5E). Like the HMG-I/Y discussed above, the analysis of snapshots from MD simulation showed that all cross-linking distance could be satisfied for the entire simulation trajectories or the selected conformation clusters (Figure S5F, 5G).

Comparing the above two different calculation methods, for restraint sampling strategy, since more distance restraints were added during the sampling process, the calculated conformation was more compact. In contrast, there were more structures that only locally form some contracted conformations during the unbiased sampling strategy, like the cluster 4,6 and 8 for HMG-I/Y, and most clusters for HMG-17 (Figure 5F, Figure S5F). Therefore, the conformations obtained by these two approaches can complement each other. Under the condition of sufficient sampling, the unbiased sampling strategy is a better and more objective choice. However, for the large protein systems, the unbiased sampling method may not be able to traverse the free energy landscape. In this case, the restraint sampling can be used as a supplement approach, especially those cross-links that cannot be satisfied in the unbiased sampling process can be used as distance restraints.

Taken together, all these results demonstrated that the CXMS could capture the different structural states of IDP objectively and reflect the intrinsic distribution of their structural states. Our study could shine a light on the of the IDP structure deciphering, which is difficult to be resolved due to the highly dynamic nature, especially in the cell. The IDP play important roles in biological processes due to their fluctuating heterogeneous conformations. However, their fuzzy structural features make the structural determination of IDPs very difficult. The IDP molecule is difficult to crystallize due to its highly dynamic feature; the disordered structure also makes the NMR spectrum less dispersed, making it difficult to resolve or analyse. In contrast, CXMS is not affected by these limitations and can directly obtain the distance information to model the IDP structures. The IDP can have different degrees of compactness and comprises transiently folded structures as functional states or interaction sites. Since the IDP lacks a stable tertiary structure, the conformations of IDP are easily affected by the surrounding environment, including the cross-linking agents. Importantly, the application of CXMS to study the structure of IDP requires careful assessment of whether these conformations are artificially induced by cross-linkers. The unbiased structural sampling could help us to identify the intrinsic movement of IDP molecules. In the two IDP molecules studied in this paper, there were indeed the intrinsic open/close movement of the whole protein, and such conformational dynamics could be accurately captured by *in vivo* CXMS.

In this paper, we demonstrate the powerful role of *in vivo* CXMS in studying intracellular protein dynamics using multiple protein systems. The dynamic structure of protein we discussed here have not been studied in cells before, and some of them have not even been studied *in vitro*. Thus, how to confirm the protein dynamic information obtained by *in vivo* CXMS is reliable? At first, out statistical analysis of nearly 5,000 proteins shows that the CXMS can unbiasedly capture the proteins of different length, types, and locations in cells. Secondly, the matching analysis of the distance between the cross-linking site and the known structure shows that most of the cross-linking pairs in those stable structural units (domains or stable secondary structures) can match the maximum arm length of the cross-linkers. Those regions that do not match the cross-linking information are mostly located in the flexible region of the proteins. Thirdly, through unbiased sampling combined with DCCM analysis, we can obtain the intrinsic motion state of the protein, and agree with cross-linking information or calculated ensemble structure from CXMS. In short, the *in vivo* CXMS can objectively capture the different conformational states of proteins, and combined with the hierarchical calculation strategy we developed, we can accurately characterize the diverse protein dynamics in cells. On the other hand, since the arm length of the cross-linker is limited, those conformational states that are not captured by the CXMS (such as the extended states) can also exist. The ensemble structure obtained by CXMS also needs further evidence from other experimental methods. At least at this stage, this structural information can help for the further functional research.

## Conclusion

The protein dynamics is closely related to their functions. The study of various conformational states changes is essential for a deeper understanding of protein functions. In this work, *in vivo* CXMS was used to represent the structure and dynamics of various proteins in the cell. With the basis of prior structural information, such as the full-length structure from PDB database or AlphaFold2 prediction, we can not only evaluate and compare the protein structures in the cell, but also characterize the conformational dynamics of different types of proteins through the distance restraints from CXMS. According to the characteristics of CXMS data, we have developed a comprehensive hierarchical strategy to illustrate the protein dynamics. A series of evaluation and calculation methods have been designed for multi-domain proteins and IDPs with different degrees of flexibility and dynamic properties. We have applied this hierarchical computational strategy to characterize dynamic structures of 3 multi-domain proteins and 2 IDP molecules. Some of the dynamic properties of these proteins have been studied *in vitro* (like CaM), while most of them have not been studied. The full-length structures of some proteins have not been resolved due to their flexibility, like hnRNP D0 and IDP molecules.

Therefore, based on the prior structural information, the CXMS enables more accurate in characterizing protein structure and dynamics in the cells. The *in vivo* CXMS with corresponding structure calculation algorithms allows intracellular structural blueprints at the proteome-wide level to explore the protein functional state in cells. In addition, *in vivo* CXMS with higher cross-linking density is required to improve its ability to decipher protein structures. New chemical cross-linking strategies, such as second-level cross-linking reaction within a shorter time scale, and location-targeted *in vivo* cross-linking to clarify the structural heterogeneity within different intracellular environments, are also needed to facilitate structural analyze with higher precise temporal and spatial resolution.

## Author contributions

Y.Z., M.L., and L.Z. conceived of the study. Z.G. and Q.Z. participated in the study design. L.Z., Y.A., and H.G. performed in vivo cross-linking experiments. B.Z. and Y.A. integrated into vivo cross-linking dataset. B.Z. performed data analysis. Z.G. performed distance calculation and structure simulation experiments. J.C. participated in the data analysis. B.Z., Z.G., Z.L., and Q.Z. wrote the manuscript. All authors read and approved the final manuscript.

## Data and materials availability

All the raw MS data have been deposited to ProteomeXchange Consortium repository with the dataset identifier PXD035828, PXD035355, PXD031128, PXD035826.

