## Supplementary Information for "A hierarchical strategy to decipher protein dynamics *in vivo* with chemical cross-linking mass spectrometry"

### 1. Experimental Procedures

#### 1.1 *In vivo* CXMS dataset

The CXMS dataset in this paper integrated our published *in vivo* cross-linking datasets from Bel-7402 by BSPNO[1], HEK293T by BSP[2], Hela by BSP[3], and Bel-7402 by BSP[4]. The detailed integrated dataset was listed in Table S1. For CXMS analysis, the cultured cells were collected and washed three times with Dulbecco's phosphate-buffered saline (PBS) before cross-linking in centrifuge tubes. The cell pellet in each group was resuspended and cross-linked in 1% (v/v) DMSO/PBS with 5 mM BSP or BSPNO at room temperature for 5 min. After cross-linking, the cell pellet was collected and resuspended in 0.2% (w/v) SDS/PBS containing a 1% (v/v) protease inhibitor cocktail to extract protein by ultrasound treatment. For click chemistry, alternative click reagent (chemistry-cleavable reagent, photo-cleavable reagent, or acid-cleavable reagent), copper (II) sulfate, THPTA, and sodium ascorbate were added. Then, 4-fold pre-cooled acetone was added to remove the excess click reagent at -20°C overnight. The supernatant was removed, and the precipitated protein pellets were dried and resuspended in 8 M urea. After reduction (8 mM DTT, RT, 1 h) and alkylation (32 mM IAA, room temperature, 30 min, dark), 50 mM ammonium bicarbonate was added to dilute urea for a final concentration of 1 M, followed by digestion with trypsin (m:m 1:50) twice at 37°C overnight. Then streptavidin agarose was added and incubated for 2 h at room temperature with slight agitation. The enriched peptides were released by 300 mM $Na_2S_2O_4$, 365 nm UV, or 10% FA. The supernatant was dried and fractionated with high-pH RPLC.

Afterward, the collected fractions of cross-linked peptides were subjected to nano LC-MS analysis on the platform of an Easy-nano LC 1200 system coupled with an Orbitrap Fusion Lumos mass spectrometer (Thermo Fisher Scientific). For CXMS data searching, the pLink 2.0 software [5] was used with parameters as follows: 20 ppm for precursor mass accuracy, 20 ppm for fragment mass accuracy, 10 ppm for precursor filter tolerance, while the results were filtered by applying a 1% FDR cut-off at the spectral level respectively for inter- and intra-cross-links. The lysine and protein N-term were set as BSP and BSPNO cross-linking sites; trypsin digestion allowed up to 3 missed cleavages, peptide length for 5-60, carbamidomethyl on [C] for fixed modification, acetyl on protein N-term and oxidation on [M] for variable modification. The data were searched against the UniprotKB human protein database.

#### 1.2 Validation of structural compatibility

Cross-links were mapped to structures from PDB (www.rcsb.org) or AlphaFold2 structure (alphafold.ebi.ac.uk). All protein structure figures were plotted using PyMol (version 2.3, Schrödinger).

### 1.3 Analysis of protein properties

The cellular component and molecular function cluster for GO analysis was performed using the DAVID database [6] (avid.ncifcrf.gov, the data was from March 2022). The protein abundance information was assigned by referring to the paxDB database [7] (pax-db.org), and the download data was further processed by the ggridges package.

### 1.4 Inter-domain dynamics calculated with CXMS restraints

The structural refinement of inter-domain dynamics against CXMS restraints was performed using Xplor-NIH [8]. The AlphaFold2 predicted structures were used as the starting structure for all three proteins. The distances between Cα atoms of cross-linked residues were used as the restraints. A square-well potential was used as the energy function, with a maximum value of 23 Å according to the arm length of the BSP and BSPNO and a minimum value of 4 Å according to the van der Walls radius of the paired atoms. There is no energy penalty when the distance between cross-linked Cα atoms is between 4-23 Å. The NTD (residues 6-76) and CTD (residues 82-149) in CaM, the two RRM domains in hnRNP A1(residues 10-88 and 106-179), and the two RRM domains in hnRNP D0 (residues 98-171 and 183-256) were treated as rigid bodies, during the structure refinement. At the same time, the intra-domain dynamic was not considered during the calculation process. The CTD was fixed to calculate CaM, and the NTD domain was grouped. For the hnRNP A1, the RRM1 domain (residues 10-88) was fixed, while the RRM2 domain (residues 106-179) was grouped. For the hnRNP D0, the RRM1 domain (residues 98-171) was fixed, and the RRM2 domain (residues 183-256) was grouped and moved around during the calculation. The residues (77-81) between NTD and CTD of CaM, the residues between two RRM domains of hnRNP A1 (89-105), and the residues between two RRMs of hnRNP D0 (172-182) all had complete torsion angle freedom during the calculations. The knowledge-based potential, including bond, angle, improper, and van der Walls repulsion, was applied in addition to the distance restraints. For CaM and hnRNP D0, only one closed conformation is required to satisfy all cross-linking restraints. However, a single conformation cannot meet all cross-linking pairs for hnRNP A1. Therefore, an alternative conformation was added for structural refinement [9]. The calculation processes for all protein systems were repeated 480 times with different random seeds. The structural ensembles satisfied all restraints, and no clashed atoms were selected for further analysis. Reweighted atomic probability maps depicting the distribution of the RRM2 domain for hnRNP A1 and hnRNP D0 were calculated in Xplor-NIH and were plotted with a threshold of 0.2[10].

### 1.5 Structure calculation of IDP molecules with *in vivo* CXMS restraints

The structure calculation with the distance restraints from CXMS was also performed using Xplor-NIH as described above. The Alphafold2 predicted conformation of

HMG-I/Y was used as the starting structure. The distances between Cα atoms of cross-linked residues were used as the restraints. The whole molecule has the full torsion freedom during the structure calculation. The calculation processes for HMG-I/Y were repeated 960 times with different random seeds. The structural ensembles satisfying all restraints, and no clashed atoms were selected for further analysis. The unbiased all atom MD simulations were performed using the AMBER 16 software package [11]. The Flexible-meccano algorithm was used to generate 5 different starting conformations with different Rg values for the HMG-I/Y and HMG-17 respectively. The protein was placed in a TIP3P explicit water box with 10 Å padding in every direction. The Particle Mesh Ewald method was used to treat the electrostatic interactions, and a 10 Å cut-off was used for non-bonded interactions [12]. A total of five independent trajectories with different random seeds were performed for the two IDP molecules. Each trajectory lasts 500 ns at 298K to produce 2500 snapshots at 200 ps intervals. The End-to-End distance ($D_{ee}$) was defined as the distance between the Cα atoms in first and last residues in the protein. The Rg and $D_{ee}$ were calculated using the CPPTRAJ module in AMBER 16. The structures were clustered using a density-based clustering algorithm [13], with a minimum of 10 points required to form a cluster and a 9 Å distance cut-off between points for creating a cluster. The centroid structure in each cluster was plotted. The dynamical cross-correlation matrix (DCCM) was calculated using CPPTRAJ module using the whole MD simulation trajectories.

### 2. Supplementary Figures and Tables

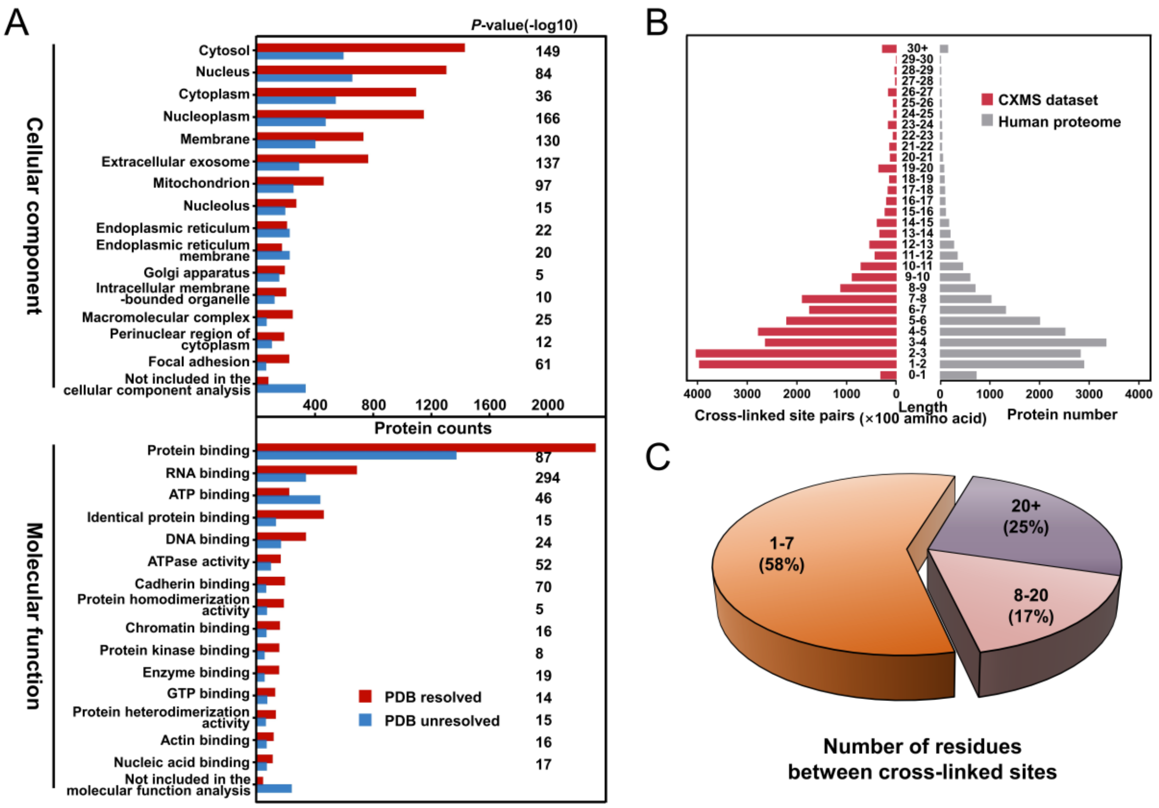

**Figure S1.** Unbiasedness of cross-linking data for structure analysis.

A) GO analysis of the cellular component and molecular function for proteins in CXMS dataset [1-4]. B) Comparison on the distribution of the identified cross-linked site pairs mapped to proteins of each sequence length and the assigned protein number from human proteome database. C) Distribution of cross-linking distance calculated by the number of residues between cross-linked sites. The number represented the cross-linking distance while the percentage was the corresponding ratio, respectively.

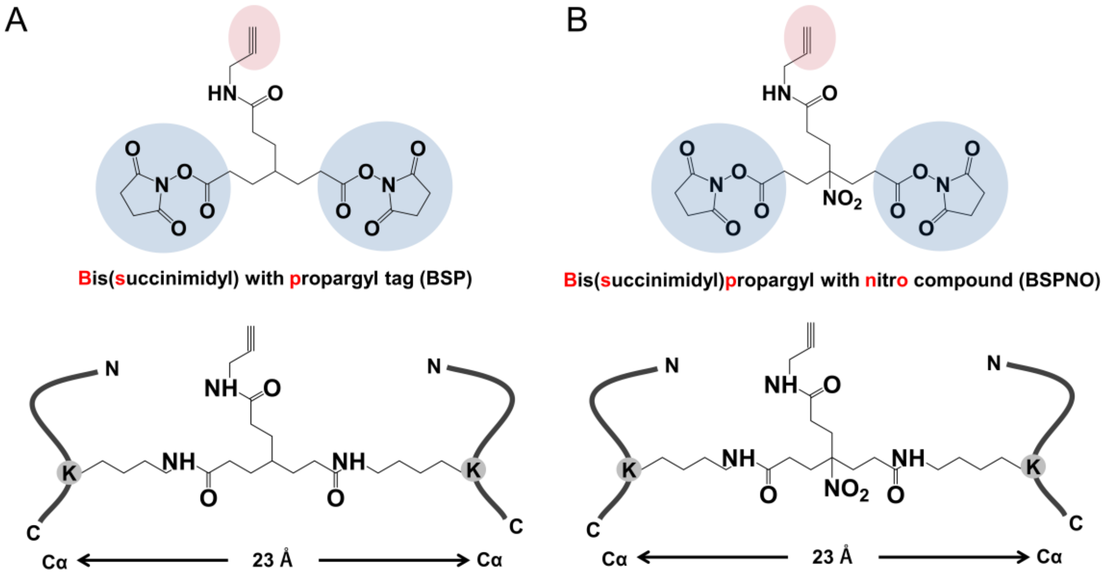

**Figure S2.** Chemical structure and maximum length of cross-linkers between Cα–Cα atoms of two cross-linked lysine residues for (A) BSP [2-4] and (B) BSPNO [1], calculated by setting all intervening dihedral angles to 180° using Chem3D 19.0 software. The maximum Cα–Cα distance restraint of the cross-linking sites was 29 Å by adding a 6 Å of protein local dynamics [14].

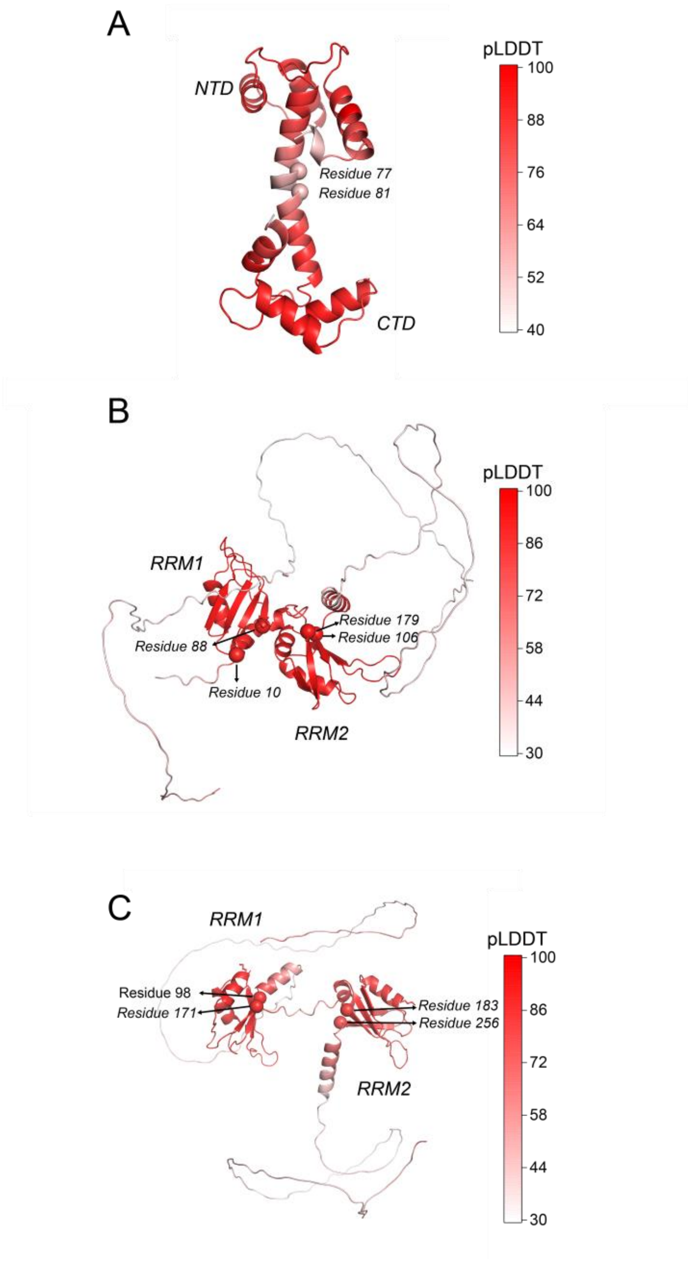

**Figure S3.** The full-length structure predicted by AlphaFold2 for calmodulin (CaM) (A), hnRNP A1 (B), and hnRNP D0 (C). The predicted local-distance difference test (pLDDT) scores of residues were indicated by colors from white to red.

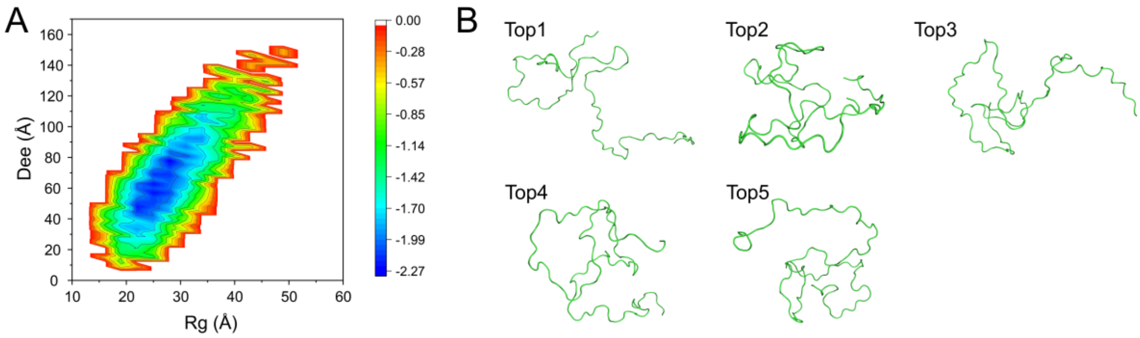

**Figure S4.** Deciphering of ensemble conformation of HMG-I/Y by the strategy of restraints sampling.

A) The 2D statistical heat map of Rg and $D_{ee}$ from the conformations satisfying all distance restraints. The more the number of structures, the small the value of the corresponding position. B) The five conformations with lowest energy scores from structure calculation with distance restraints from *in vivo* CXMS.

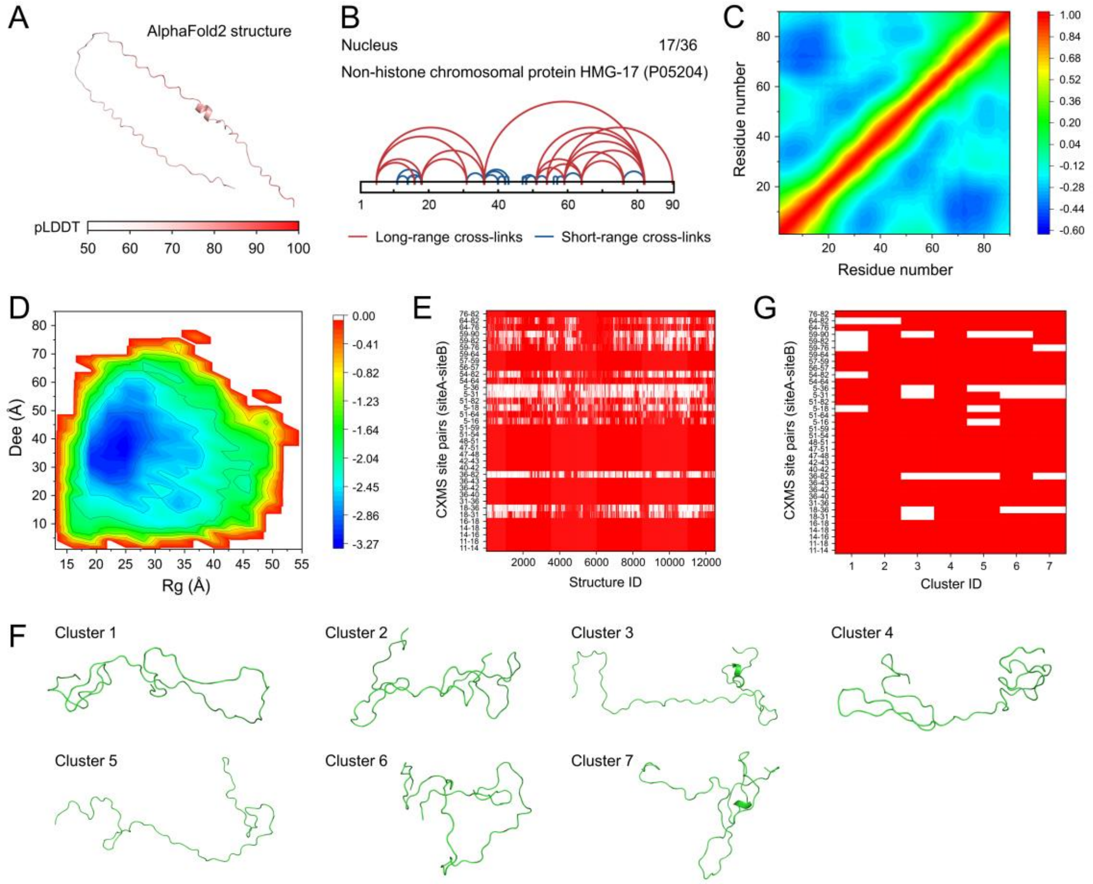

**Figure S5.** Deciphering of ensemble structure of HMG-17 by the strategy of unbiasedly sampling and reweighting.

A) The full-length structure of HMG-17 predicted by AlphaFold2. The pLDDT scores of residues were indicated by colors from white to red. B) All identified cross-links were mapped onto the 2D structure of HMG-17. The short- (number of residues between cross-linked sites less than 8) and long- range cross-linking information were colored in blue and red, respectively. The number of long-range cross-links and the total number of cross-linked pairs were indicated and separated by “/”. C) The DCCM analysis of HMG-17. The correlation coefficient related with residue related movement were colored from blue to red. The stronger the negative correlation, the small the value of the corresponding position. D) The 2D statistical heat map of Rg and $D_{ee}$ for the MD simulation trajectories of HMG-17. The more the number of structures, the small the value of the corresponding position. E) The matching between each snapshot from MD simulation and corresponding CXMS information. The cross-linking distance satisfying or not was shown in red and white, respectively. F) The representative structure of seven clusters from unbiased all atom MD simulations. G) The matching between seven structure clusters and corresponding CXMS information. The cross-linking distance satisfying or not was shown in red and white, respectively.

**Table S2.** Cross-linked sites in domains for CaM.

| **Cross-link** | **Spectra** | **$C_\alpha$-$C_\alpha$ (Å)** | **Compatible[a]** | **Remark** |
|---|---|---|---|---|
| $Lys^{14}$-$Lys^{95}$ | 1 | 39.04 | No | inter-domain |
| $Lys^{22}$-$Lys^{95}$ | 23 | 47.08 | No | inter-domain |
| $Lys^{22}$-$Lys^{31}$ | 189 | 9.79 | Yes | NTD |
| $Lys^{14}$-$Lys^{22}$ | 19 | 12.81 | Yes | NTD |
| $Lys^{76}$-$Lys^{78}$ | 56 | 5.40 | Yes | Loop[b] |

[a] Whether the corresponding distance was less than the arm length of cross-linker
[b] At least one cross-linked site was located at the loop region between domains

**Table S3.** Cross-linked sites in domains for hnRNP A1.

| **Cross-link** | **Spectra** | **$C_\alpha$-$C_\alpha$ (Å)** | **Compatible[a]** | **Remark** |
|---|---|---|---|---|
| $Lys^{15}$-$Lys^{179}$ | 6 | 18.92 | Yes | inter-domain |
| $Lys^{78}$-$Lys^{106}$ | 23 | 25.63 | Yes | inter-domain |
| $Lys^{78}$-$Lys^{179}$ | 6 | 26.49 | Yes | inter-domain |
| $Lys^{87}$-$Lys^{106}$ | 22 | 14.91 | Yes | inter-domain |
| $Lys^{87}$-$Lys^{166}$ | 2 | 17.51 | Yes | inter-domain |
| $Lys^{87}$-$Lys^{179}$ | 6 | 13.58 | Yes | inter-domain |
| $Lys^{52}$-$Lys^{113}$ | 3 | 52.63 | No | inter-domain |
| $Lys^{52}$-$Lys^{179}$ | 3 | 33.57 | No | inter-domain |
| $Lys^{78}$-$Lys^{113}$ | 5 | 44.21 | No | inter-domain |
| $Lys^{78}$-$Lys^{166}$ | 9 | 31.68 | No | inter-domain |
| $Lys^{87}$-$Lys^{113}$ | 1 | 31.50 | No | inter-domain |
| $Lys^{15}$-$Lys^{52}$ | 1 | 23.03 | Yes | RRM1 |
| $Lys^{15}$-$Lys^{78}$ | 8 | 18.32 | Yes | RRM1 |
| $Lys^{15}$-$Lys^{87}$ | 13 | 8.82 | Yes | RRM1 |
| $Lys^{52}$-$Lys^{78}$ | 4 | 22.66 | Yes | RRM1 |
| $Lys^{78}$-$Lys^{87}$ | 10 | 14.27 | Yes | RRM1 |
| $Lys^{106}$-$Lys^{113}$ | 6 | 19.99 | Yes | RRM2 |
| $Lys^{113}$-$Lys^{179}$ | 6 | 19.93 | Yes | RRM2 |
| $Lys^{161}$-$Lys^{166}$ | 13 | 10.57 | Yes | RRM2 |
| $Lys^{161}$-$Lys^{179}$ | 7 | 10.83 | Yes | RRM2 |
| $Lys^{166}$-$Lys^{179}$ | 33 | 10.40 | Yes | RRM2 |
| $Lys^{15}$-$Lys^{105}$ | 22 | 19.82 | Yes | loop[b] |
| $Lys^{78}$-$Lys^{105}$ | 25 | 21.90 | Yes | loop[b] |
| $Lys^{87}$-$Lys^{105}$ | 67 | 12.27 | Yes | loop[b] |
| $Lys^{52}$-$Lys^{105}$ | 6 | 34.43 | No | loop[b] |
| $Lys^{105}$-$Lys^{106}$ | 2 | 3.86 | Yes | loop[b] |
| $Lys^{105}$-$Lys^{113}$ | 23 | 23.50 | Yes | loop[b] |
| $Lys^{105}$-$Lys^{179}$ | 26 | 8.22 | Yes | loop[b] |

[a] Whether the corresponding distance was less than the arm length of cross-linker
[b] At least one cross-linked site was located at the loop region between domains

**Table S4.** Cross-linked sites in domains for hnRNP D0.

| Cross-link | Spectra | $C_\alpha$-$C_\alpha$ (Å) | Compatible[a] | Remark |
|---|---|---|---|---|
| Lys$^{146}$-Lys$^{197}$ | 3 | 49.87 | No | inter-domian |
| Lys$^{110}$-Lys$^{111}$ | 36 | 3.85 | Yes | RRM1 |
| Lys$^{111}$-Lys$^{114}$ | 33 | 5.01 | Yes | RRM1 |
| Lys$^{161}$-Lys$^{165}$ | 4 | 5.48 | Yes | RRM1 |
| Lys$^{110}$-Lys$^{114}$ | 28 | 6.26 | Yes | RRM1 |
| Lys$^{114}$-Lys$^{119}$ | 2 | 8.91 | Yes | RRM1 |
| Lys$^{119}$-Lys$^{153}$ | 1 | 9.54 | Yes | RRM1 |
| Lys$^{153}$-Lys$^{158}$ | 1 | 10.39 | Yes | RRM1 |
| Lys$^{158}$-Lys$^{161}$ | 10 | 10.42 | Yes | RRM1 |
| Lys$^{98}$-Lys$^{129}$ | 1 | 14.61 | Yes | RRM1 |
| Lys$^{119}$-Lys$^{129}$ | 1 | 22.02 | Yes | RRM1 |
| Lys$^{242}$-Lys$^{243}$ | 32 | 3.86 | Yes | RRM2 |
| Lys$^{237}$-Lys$^{238}$ | 2 | 3.88 | Yes | RRM2 |
| Lys$^{218}$-Lys$^{221}$ | 44 | 8.17 | Yes | RRM2 |
| Lys$^{243}$-Lys$^{255}$ | 16 | 8.72 | Yes | RRM2 |
| Lys$^{243}$-Lys$^{251}$ | 9 | 10.21 | Yes | RRM2 |
| Lys$^{242}$-Lys$^{251}$ | 19 | 11.70 | Yes | RRM2 |
| Lys$^{251}$-Lys$^{255}$ | 4 | 13.52 | Yes | RRM2 |
| Lys$^{170}$-Lys$^{173}$ | 2 | 8.23 | Yes | loop[b] |
| Lys$^{129}$-Lys$^{178}$ | 8 | 18.84 | Yes | loop[b] |
| Lys$^{178}$-Lys$^{237}$ | 6 | 13.55 | Yes | loop[b] |
| Lys$^{176}$-Lys$^{178}$ | 24 | 5.09 | Yes | loop[b] |
| Lys$^{178}$-Lys$^{182}$ | 1 | 10.55 | Yes | loop[b] |
| Lys$^{173}$-Lys$^{178}$ | 4 | 13.00 | Yes | loop[b] |

[a] Whether the corresponding distance was less than the arm length of cross-linker

[b] At least one cross-linked site was located at the loop region between domains

**Table S5** Cross-linked sites for HMG-I/Y.

| Cross-link | Spectra | $C_\alpha$-$C_\alpha$ (Å) | Compatible[a] |
|---|---|---|---|
| $Lys^{88}$-$Lys^{89}$ | 1 | 3.814 | Yes |
| $Lys^{65}$-$Lys^{67}$ | 5 | 6.06 | Yes |
| $Lys^{71}$-$Lys^{74}$ | 17 | 8.002 | Yes |
| $Lys^{15}$-$Lys^{18}$ | 112 | 8.255 | Yes |
| $Lys^{18}$-$Lys^{23}$ | 2 | 11.944 | Yes |
| $Lys^{15}$-$Lys^{23}$ | 3 | 17.801 | Yes |
| $Lys^{67}$-$Lys^{74}$ | 4 | 17.875 | Yes |
| $Lys^{74}$-$Lys^{88}$ | 1 | 19.612 | Yes |
| $Lys^{7}$-$Lys^{15}$ | 14 | 22.817 | Yes |
| $Lys^{74}$-$Lys^{89}$ | 10 | 22.852 | Yes |
| $Lys^{65}$-$Lys^{74}$ | 17 | 23.842 | Yes |
| $Lys^{15}$-$Lys^{31}$ | 7 | 24.954 | Yes |
| $Lys^{7}$-$Lys^{18}$ | 52 | 28.441 | Yes |
| $Lys^{62}$-$Lys^{74}$ | 1 | 30.163 | No |
| $Lys^{31}$-$Lys^{46}$ | 28 | 31.511 | No |
| $Lys^{7}$-$Lys^{23}$ | 1 | 31.573 | No |
| $Lys^{7}$-$Lys^{31}$ | 37 | 40.969 | No |
| $Lys^{46}$-$Lys^{67}$ | 2 | 43.229 | No |
| $Lys^{15}$-$Lys^{46}$ | 5 | 46.158 | No |
| $Lys^{7}$-$Lys^{62}$ | 1 | 50.306 | No |
| $Lys^{46}$-$Lys^{74}$ | 30 | 50.66 | No |
| $Lys^{7}$-$Lys^{67}$ | 4 | 60.264 | No |
| $Lys^{7}$-$Lys^{55}$ | 1 | 61.504 | No |
| $Lys^{18}$-$Lys^{74}$ | 1 | 62.845 | No |
| $Lys^{7}$-$Lys^{46}$ | 39 | 67.2 | No |
| $Lys^{15}$-$Lys^{74}$ | 2 | 68.369 | No |
| $Lys^{7}$-$Lys^{74}$ | 36 | 77.315 | No |

[a] Whether the corresponding distance was less than the arm length of cross-linker

**Table S6.** Cross-linked sites for HMG-17.

| **Cross-link** | **Spectra** | **$C_\alpha$-$C_\alpha$ (Å)** | **Compatible[a]** |
|---|---|---|---|
| Lys$^{47}$-Lys$^{48}$ | 14 | 3.74 | Yes |
| Lys$^{56}$-Lys$^{57}$ | 11 | 3.76 | Yes |
| Lys$^{42}$-Lys$^{43}$ | 5 | 3.82 | Yes |
| Lys$^{14}$-Lys$^{16}$ | 4 | 5.51 | Yes |
| Lys$^{16}$-Lys$^{18}$ | 20 | 5.65 | Yes |
| Lys$^{40}$-Lys$^{42}$ | 33 | 5.90 | Yes |
| Lys$^{57}$-Lys$^{59}$ | 11 | 6.79 | Yes |
| Lys$^{48}$-Lys$^{51}$ | 4 | 8.90 | Yes |
| Lys$^{11}$-Lys$^{14}$ | 20 | 8.96 | Yes |
| Lys$^{51}$-Lys$^{54}$ | 2 | 9.18 | Yes |
| Lys$^{47}$-Lys$^{51}$ | 1 | 10.62 | Yes |
| Lys$^{14}$-Lys$^{18}$ | 10 | 11.02 | Yes |
| Lys$^{36}$-Lys$^{40}$ | 14 | 11.17 | Yes |
| Lys$^{59}$-Lys$^{64}$ | 4 | 13.10 | Yes |
| Lys$^{31}$-Lys$^{36}$ | 4 | 14.27 | Yes |
| Lys$^{36}$-Lys$^{42}$ | 8 | 15.87 | Yes |
| Lys$^{76}$-Lys$^{82}$ | 49 | 17.32 | Yes |
| Lys$^{36}$-Lys$^{43}$ | 25 | 19.31 | Yes |
| Lys$^{11}$-Lys$^{18}$ | 9 | 19.71 | Yes |
| Lys$^{18}$-Lys$^{31}$ | 6 | 20.86 | Yes |
| Lys$^{51}$-Lys$^{59}$ | 3 | 21.71 | Yes |
| Lys$^{54}$-Lys$^{64}$ | 2 | 25.76 | Yes |
| Lys$^{51}$-Lys$^{64}$ | 2 | 29.77 | No |
| Lys$^{5}$-Lys$^{16}$ | 8 | 30.50 | No |
| Lys$^{64}$-Lys$^{76}$ | 2 | 31.81 | No |
| Lys$^{18}$-Lys$^{36}$ | 4 | 34.45 | No |
| Lys$^{5}$-Lys$^{18}$ | 6 | 34.89 | No |
| Lys$^{36}$-Lys$^{82}$ | 3 | 41.95 | No |
| Lys$^{59}$-Lys$^{76}$ | 3 | 44.67 | No |
| Lys$^{64}$-Lys$^{82}$ | 8 | 49.01 | No |
| Lys$^{5}$-Lys$^{31}$ | 1 | 55.35 | No |
| Lys$^{59}$-Lys$^{82}$ | 2 | 61.88 | No |
| Lys$^{5}$-Lys$^{36}$ | 4 | 68.49 | No |
| Lys$^{54}$-Lys$^{82}$ | 2 | 71.67 | No |
| Lys$^{51}$-Lys$^{82}$ | 7 | 71.68 | No |
| Lys$^{59}$-Lys$^{90}$ | 6 | 83.29 | No |

[a] Whether the corresponding distance was less than the arm length of cross-linker
.
